# Severe murine schistosomiasis results from disruption of the modulation of anti-egg CD4 T cell response by immunodominance of a single egg epitope

**DOI:** 10.1101/2020.05.22.110791

**Authors:** Eduardo Finger, Thaissa Melo Galante Coimbra, Alessandra Finardi de Souza

**Author notes:** Corresponding Author: Eduardo Finger MD PhD ^A, B^, Rua 13 de Maio 1954 cj 114, São Paulo – SP, CEP 01327-002 - Brazil.

## Abstract

This study exploits the consistent correlation between immunodominance of the major egg antigen Sm-p40_234-246_, a robust Th1/Th17 anti-egg CD4 response and severe liver immunopathology in experimental murine schistosomiasis as an experimental platform to analyze how different degrees of immunodominance affect CD4 modulation and disease outcome. The results show that strong immunodominance of a restricted egg epitope repertoire skews CD4 modulation towards a pathogenic Th1/Th17 pro-inflammatory response and that neutralizing this immunodominance generates an opposite and restorative effect. These results identify immunodominance as an important pathogenic component that influences CD4 modulation in experimental murine schistosomiasis and can be manipulated to treat this and maybe other CD4 mediated diseases.

**Summary:** Antigen informed CD4 modulation determines how efficiently the immune system neutralizes a threat; however, this process and its components are not fully comprehended. This study analyzes immunodominance as one component able to disrupt CD4 modulation and turn pathogenic an otherwise healthy immune response.

## Introduction

When facing a challenge, the adaptive immune system seeks for the best strategy to neutralize it with the least possible collateral damage[1] and the information required for this task mostly derives from the interaction between APCs and CD4s, the former providing epitopes, cytokines and costimulation, the latter processing them into threat specific strategies e.g. Th1 responses for intracellular organisms, Th2 against helminths or toxins or Th17 for extracellular organisms. If the strategy fits that challenge and is properly executed, the threat is neutralized, otherwise, the result may be prolonged illness and immunopathology. The process of adjusting the strategy to the challenge is called modulation and, although not entirely understood, it involves the dynamic interaction of at least 6 components: innate immunity signals[2], epitope concentration[3], epitope affinity for the MHC[4], TCR affinity for the MHC/epitope ligand[5], costimulation[6, 7] and the local immune environment[8].

Murine schistosomiasis egg-induced liver immunopathology (SELI) is a useful and consistent model to study modulation[9] and has helped elucidate the role of several of its components [10, 11]. In this model, infection with *S.mansoni* initially induces a mild Th1 response until 4 wpi, when mature schistosomes start laying eggs, some of which get trapped in the portal spaces of the liver where they elicit a CD4 mediated granulomatous response[12]. Infected C57BL/6 (H-2B) and Balb/c (H-2D) mice represent low pathology strains because they respond to these eggs by modulating that initial Th1 response to a predominantly Th2 anti-egg response conducive to small, well delimited perioval granulomas and minor liver damage. Conversely, CBA/J and C3H/HeN mice (both H-2K) represent high pathology strains because upon contact with eggs, they boost the initial Th1 response into a stronger, polarized Th1/Th17 response that produces large, infiltrative, highly inflammatory perioval granulomas, disseminated liver damage and severe disease.

Analysis of the CD4 reaction against purified egg antigens show that high and low pathology strains segregate according to their response the major Sm-p40 egg antigen or, specifically, its immunodominant epitope, Sm-p40_234-246_. While CD4s from high pathology strains respond to Sm-p40 with the same strong Th1/Th17 polarization observed in experimental infection, CD4s from low pathology ones remain unresponsive[13, 14]. This observation suggests that anti-Sm-p40 CD4 reactivity may originate and be pivotal for the pro-Th1/Th17 polarization that precedes the development of high pathology[15].

Sm-p40 is a 354 amino acid glycoprotein that comprises 10% of the all the egg’s proteins[16]. At the MHCII epitope presentation level, I-A^k^ restricts Sm-p40 to 2 subdominant epitopes (Sm-p40_179-208_ and Sm-p40_279-308_) and a major dominant one (Sm-p40_234-246_)[14], the latter being the target of the Vα11.3β8 CD4 clonotype that comprises most of the anti-egg CD4 repertoire identified in both high pathology strains but not low pathology ones[17]. I-A^b^ restricts Sm-p40 to 5 epitopes and I-A^d^, to 18, none of them majorly dominant (Table 1). However, could the outcome of an intricate disease like schistosomiasis be determined by CD4 responsivity against a single epitope?

**Table 1.** Number of Sm-p40 epitopes predicted to be presented by different MHCII haplotypes per the RANKPEP algorithm

| Haplotype | Binding threshold<sup>a</sup> | Number of Sm-p40 epitopes presented<sup>b</sup> |
| --- | --- | --- |
| I-A<sup>k</sup> | 14.17 | 2 |
| I-A<sup>b</sup> | 9.52 | 5 |
| I-A<sup>d</sup> | 7.1 | 18 |
| I-A<sup>s</sup> | 12.65 | 1 |

<sup>a</sup> Minimal score required for an epitope to be able to bind to the MHC molecule and be presented

One possible explanation is ligand density (LD), a concept proposed by Bottomly and Murray[4, 18–20] wherein, as a competitive process, MHCII epitope loading and presentation hinge on that epitope concentration and affinity for the MHCII, thus, in a situation where none prevail, many are presented in a diverse but low LD conducive to a Th2 response. However, if one epitope outcompetes others in either or both these attributes, it achieves a high LD that elicits a Th1 response[19, 21]. As such, Sm-p40 epitomizes the perfect storm against effective modulation of the anti-egg CD4 response in H-2K strains, for, not only it is the most abundant antigen within eggs, but its dominant epitope exhibits uniquely high affinity for I-A^k^, which, itself, is a more stringent than average MHCII haplotype[14, 22]. Thus, altogether, Sm-p40_234-246_ dominates MHCII antigen presentation on H-2K APCs with such LD that the result is a pro-Th1/Th17 stimulus whose strength overcomes the fiercest modulation effort. This would explain why, despite secreting severalfold more Th2 cytokines than low pathology strains, the anti-egg CD4 response of high pathology ones still degenerates into a severe liver immunopathology.

This study analyzes the effect of restrictive immunodominance^*^ (RI) in CD4 modulation and its immunopathogenic outcome.

## Results

### A. Sm-p40_234-246_ RI neutralization prevents the development of severe SELI in CBA mice

If Sm-p40_234-246_ RI is critical for the development of severe SELI, its neutralization should allow for adequate modulation of the anti-egg CD4 response and, thus, prevent it.

To test this hypothesis, we immunized infected CBA mice (Fig. 1) with a cocktail of 4 synthetic epitopes selected for being expressed within egg proteins, and possessing an affinity for I-A^k^ that rival that of Sm-p40234-246(Table 1), such that now, APCs can present 5 epitopes instead of one, matching the number of Sm-p40 epitopes presented by APCs in the low pathology C57BL/6 mouse (Table 2).

**Fig. 1.**
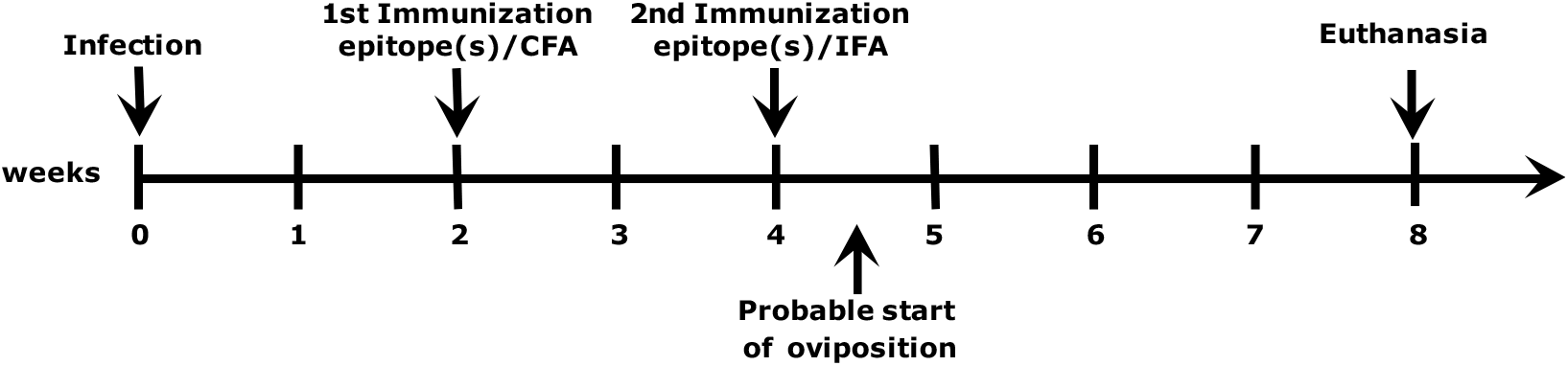
Schematic illustration of the experimental protocol for the neutralization of Sm-p40_234-246_ RI in CBA mice or induction of CPP_1380-1397_ RI in C57BL/6 mice.

**Table 2.**
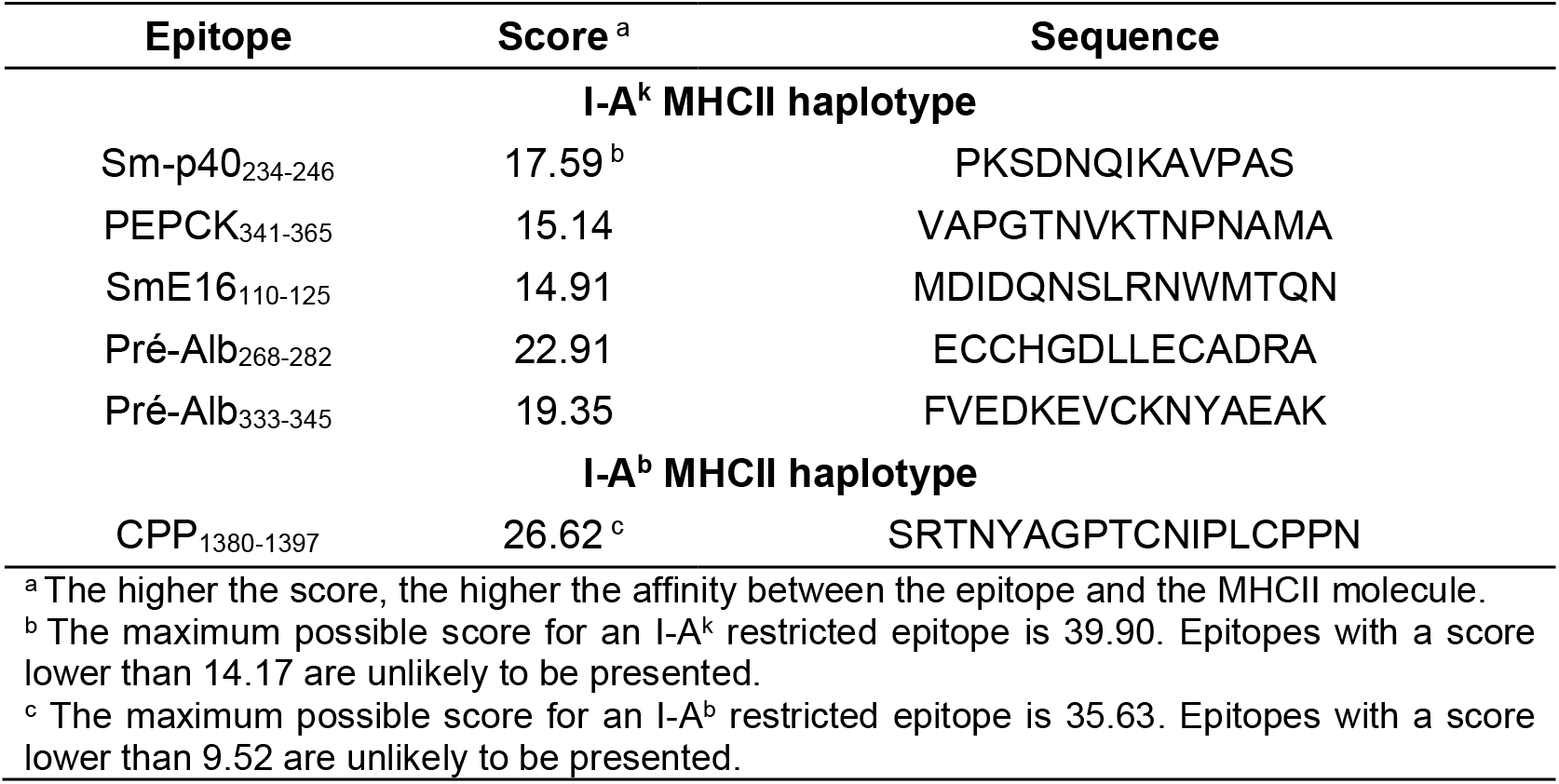
MHCII/Epitope affinity as predicted by the RANKPEP algorithm

| Epitope | Score <sup>a</sup> | Sequence |
| --- | --- | --- |
| **I-A<sup>k</sup> MHCII haplotype** |  |  |
| Sm-p40<sub>234-246</sub> | 17.59 <sup>b</sup> | PKSDNQIKAVPAS |
| PEPCK<sub>341-365</sub> | 15.14 | VAPGTNVKTNPNAMA |
| SmE16<sub>110-125</sub> | 14.91 | MDIDQNSLRNWMTQN |
| Pré-Alb<sub>268-282</sub> | 22.91 | ECCHGDLLECADRA |
| Pré-Alb<sub>333-345</sub> | 19.35 | FVEDKEVCKNYAEAK |
| **I-A<sup>b</sup> MHCII haplotype** |  |  |
| CPP<sub>1380-1397</sub> | 26.62 <sup>c</sup> | SRTNYAGPTCNIPLCPPN |

<sup>a</sup> The higher the score, the higher the affinity between the epitope and the MHCII molecule.
<sup>b</sup> The maximum possible score for an I-A<sup>k</sup> restricted epitope is 39.90. Epitopes with a score lower than 14.17 are unlikely to be presented.
<sup>c</sup> The maximum possible score for an I-A<sup>b</sup> restricted epitope is 35.63. Epitopes with a score lower than 9.52 are unlikely to be presented.

Morphometric analysis of liver granulomas shows that RI neutralization prevents the development of severe SELI in CBA mice in an almost dose dependent way, to the point where CBA and C57BL/6 SELI become indistinguishable (Fig. 2, 5). The same protocol using a mock cocktail of 4 random peptides with no affinity for I-A^k^ did not affect CBA SELI (data not shown).

**Fig. 2.**
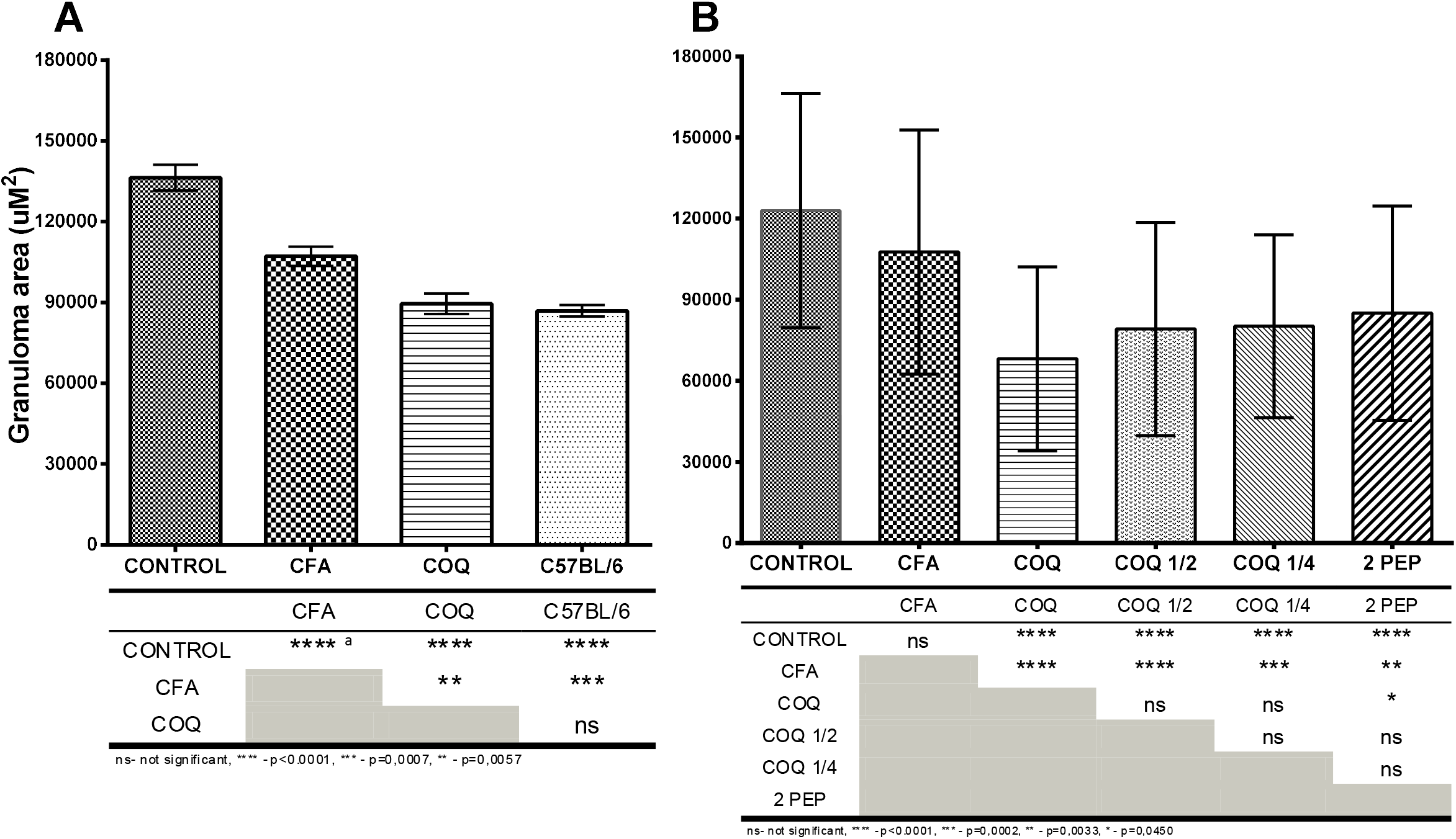
Morphometric analysis of egg-induced liver granulomas showing that Sm-p40_234-246_ RI neutralization reduces the severity of SELI in Schistosome infected female CBA mice. **A** – Experiment comparing the SELI of untreated CBA mice (CONTROL), mice immunized solely with adjuvant (CFA), mice that underwent Sm-p40_234-246_ RI neutralization strategy (COQ) and untreated low pathology C57BL/6 mice. **B** – Experiment comparing the effect of full Sm-p40_234-246_ RI neutralizing strategy (COQ), with half the dose of the epitope cocktail (COQ ½), a quarter of the dose (COQ ¼) or a different composition of the cocktail that contained 2 epitopes instead of the regular 4 (2 PEP). The table below the graphs shows the statistical differences between the test groups. Results are representative of 6 different experiments. No test group had less than 3 mice or 78 granulomas. Use of a mock cocktail without affinity for I-Ak produced no change in CBA SELI. a – In this experiment, the SELI of group CFA differed significantly from group CONTROL. This is not the typical outcome for this group and is compensated by the fact that group COQ also differs significantly from group CFA.

### B. Non-H-2K strains subject to Sm-p40 RI develop severe SELI

If Sm-p40 RI is responsible for the polarization that aggravates SELI, that should happen whenever it occurs, irrespective of the strain.

To verify this hypothesis, we analyzed Sm-p40 restriction patterns in several MHCII haplotypes and identified that Sm-p40 restriction in the I-A^s^ haplotype is even stricter than the I-A^k^ haplotype (Table 2), and therefore, predicted that, infected with *S. mansoni*, H-2S SJL mice should develop CBA-like SELI.

Morphometric results show that, as predicted, infected SJL mice develop as severe a SELI as CBA mice (Fig. 3, 5), with a non-significant trend towards severer disease.

**Fig. 3.**
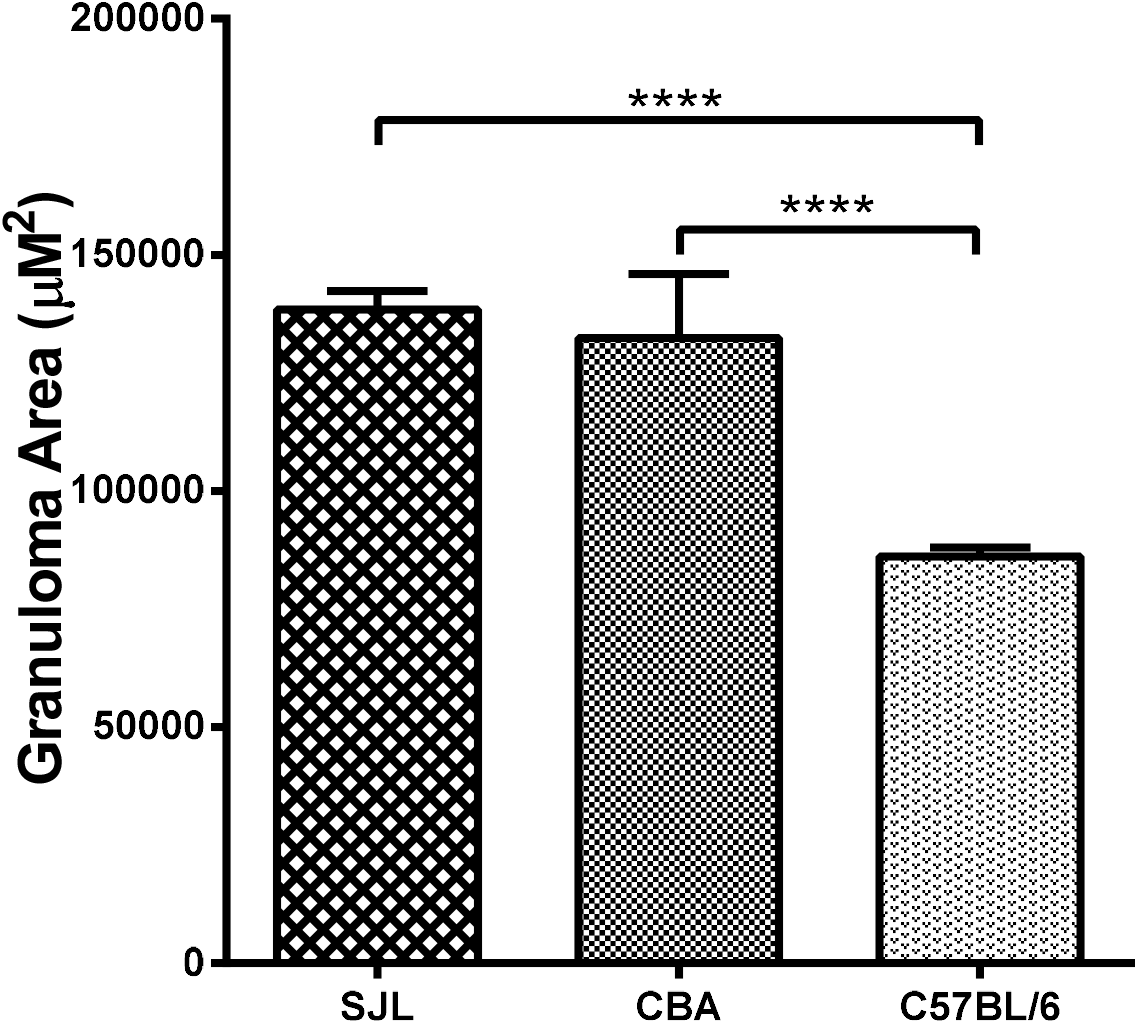
Morphometric analysis of egg-induced liver granulomas from schistosome infected SJL, CBA and C57BL/6 mice showing that in a different MHCII haplotype, severe SELI still correlates with restrictive immunodominance of Sm-p40. Results are representative of 4 different experiments. No test group had less than 3 mice or 176 granulomas.

### C. Experimentally induced RI leads to severe SELI in C57BL/6 mice

To test whether a single epitope can turn pathogenic an otherwise adequate anti-egg CD4 response, in a reversal of the Sm-p40 RI neutralization assay, we scrutinized the *S.mansoni* genome to identify the highest ranked epitope in terms of affinity for I-A^b^, that is expressed within an egg protein(Table 1), and used it to immunize C57BL/6 mice(Fig. 1) and verify the effect of experimentally induced RI on SELI.

Morphometric analysis shows that sensitization with this RI inducing epitope suffices to significantly aggravate SELI in C57BL/6 mice (Fig. 4, 5)

**Fig. 4.**
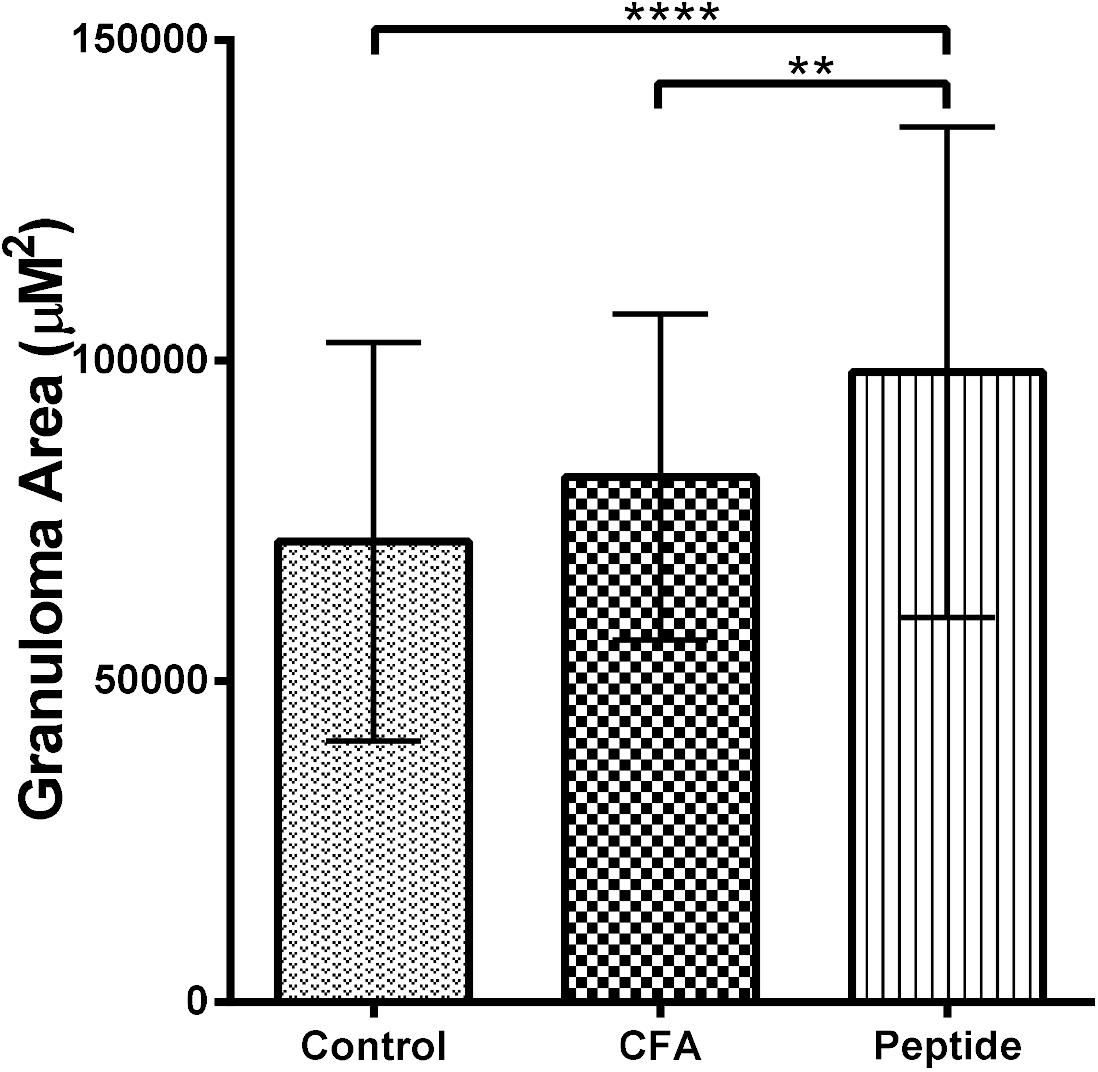
Morphometric analysis of egg-induced liver granulomas from schistosome infected C57BL/6 mice untreated (Control), immunized solely with adjuvant (CFA), or adjuvant plus CPP1380-1397 (Peptide), a synthetic epitope capable of inducing restrictive immunodominance in I-Ab mice, showing that severe SELI correlated with induction of RI by a single epitope.

**Fig. 5.**
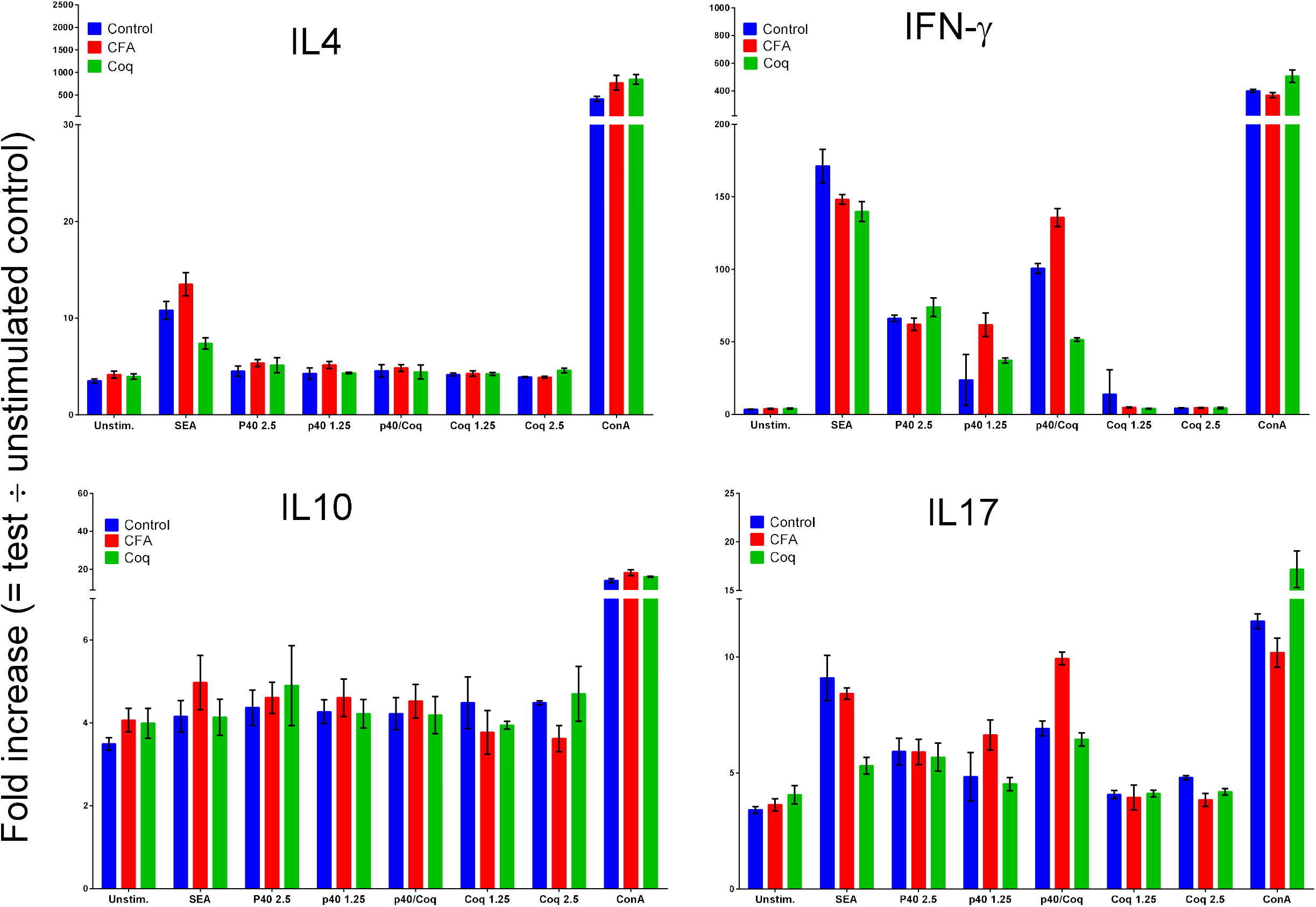
Analysis of the ex vivo production of IL4, IL10, IFN-γ and IL17 by CD4s purified from the mesenteric lymph nodes of CBA mice 8wpi, subjected to the standard RI neutralization protocols, stimulated by SEA at 25 μg/ml(SEA), Sm-p40(p40) at 2.5μM and 1.25 μM, RI neutralization cocktail at 2.5μM and 1.25μM(Coq) and both(p40/coq) at 1.25 μM each(p40/Coq) and concanavalin A at 75μg/ml(ConA). As previously described, Sm-p40 does not elicit IL4 or IL10 in CBA mice but elicits a strong IFN-γ and IL17 production, curtailed by RI neutralization (Coq) when compared to mice who received only adjuvant (CFA).

**Fig. 6.**
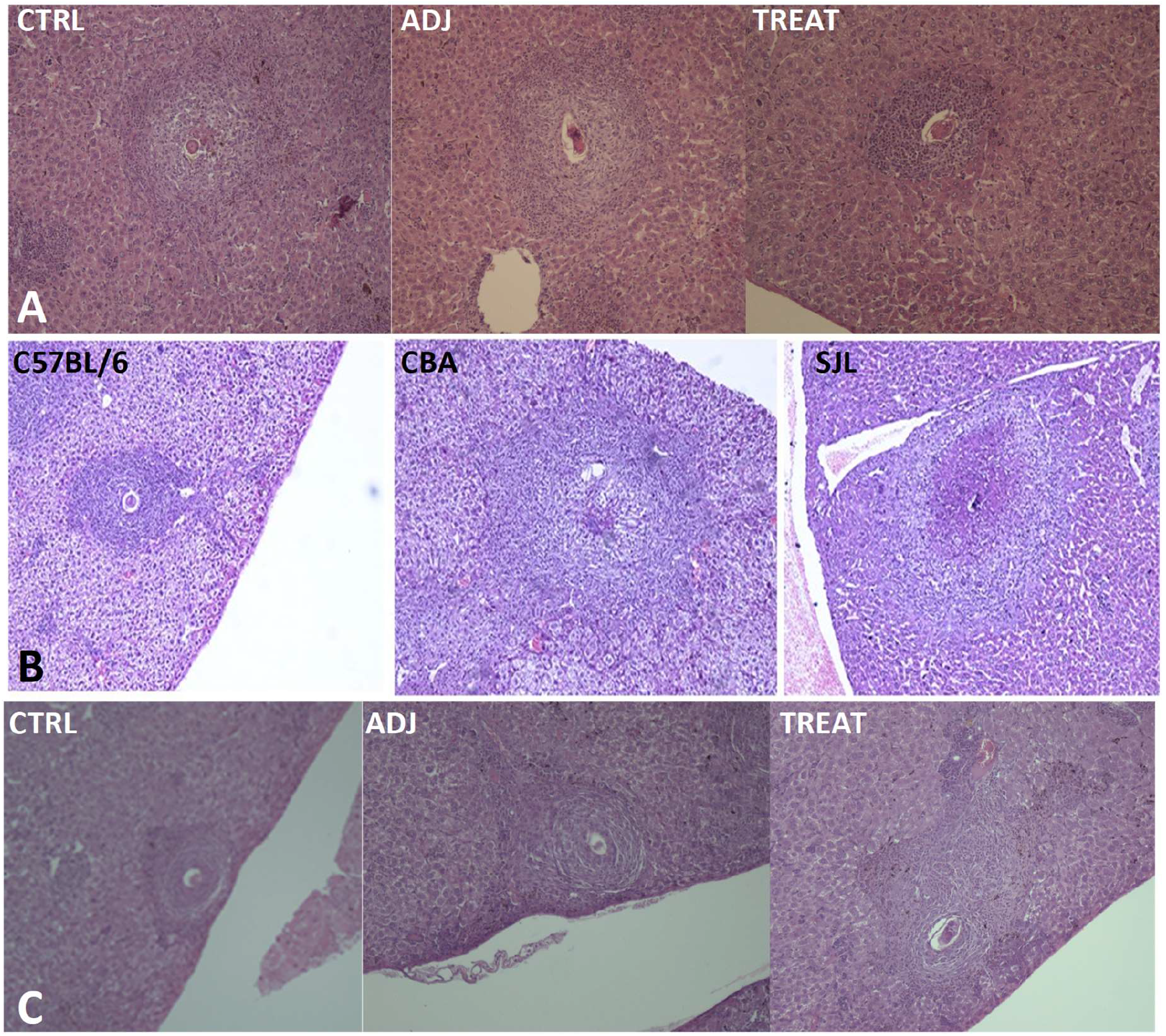
Comparative liver immunopathology of the three experimental groups analyzed. A – Sm-p40+ RI neutralization experiment showing representative granulomas from the untreated CBA controls (CTRL), CBA mice treated with adjuvant only (ADJ) and CBA mice which underwent RI neutralization strategy (TREAT). B – Experiment showing representative granulomas in different strains subject (CBA and SJL) or not (C57BL/6) to Sm-p40 RI. C – Experiment showing representative granulomas from untreated C57BL/6 mice (CTRL), C57BL/6 mice treated with adjuvant alone (ADJ) or C57BL/6 mice treated with the RI inducing peptide CPP_1380-1397_.

### D. Sm-p40_234-246_ RI neutralization reduces cytokine polarization in infected CBA mice

If Sm-p40_234-246_ RI causes the strong pro-Th1/Th17 stimulus observed in severe schistosomiasis, its neutralization should reduce the production of its signature cytokines, respectively, interferon gamma (IFN-γ) and IL-17.

To verify this hypothesis, we analyzed, the *ex vivo* production of IL-4, IL-17, IFN-γ and IL-10 in CD4s purified from mesenteric lymph nodes of 8wpi CBA mice, undergoing, or not, RI neutralization protocol.

The results show that mice undergoing RI neutralization protocol consistently display lower levels of IFN-γ and IL-17 against Sm-p40, compared to mice immunized with adjuvant alone. As previously mentioned, Sm-p40 does not elicit Th2 cytokines in CBA mice[9].

### E. Is RI relevant for human CD4 mediated diseases?

To verify whether we can identify a role for RI within human CD4 mediated diseases, we performed a RANKPEP restriction analysis of three human diseases selected for combining a dominant CD4 mediated pathogenesis, with a strong correlation with a specific MHCII haplotype and a known, or strongly suspected, eliciting antigen: Lyme disease (OspA/DRB1*0401)[23], celiac disease (gamma-gliadin/DQ2.5)[24] and multiple sclerosis (myelin/DRB1*1501/02)[25] (Table 3).

**Table 3.** MHCII specific restriction analysis of antigens correlated with CD4 mediated autoimmune diseases using the RANKPEP algorithm

| CD4 mediated disease | Eliciting Antigen | MHCII Haplotype Restriction | Number of competitive Epitopes | MHCII affinity score of fittest epitopes relative to maximum score<sup>c</sup> | MHCII affinity ratio between the 1<sup>st</sup> and 2<sup>nd</sup> fittest epitopes<sup>d</sup> |
| --- | --- | --- | --- | --- | --- |
| Lyme Disease | OspA | DRB1*0401 | 10<sup>a</sup> | 52,52% | 2,50 |
| Celiac disease | gliadin | DQ2.5 | 1 | 47,78% | 1.28 |
| Multiple sclerosis | myelin<sup>e</sup> | DRB1*1501 | 10<sup>b</sup> | 96,81% | 2.34 |
|  |  | DRB1*1502 | 3 | 81,03% | 6.27 |

<sup>a</sup> Although RANKPEP predicts 10 possible epitopes, several of them are overlaps of the same epitope, therefore, there are only 5 unique epitopes. <sup>b</sup> Although RANKPEP predicts 10 possible epitopes, several of them are overlaps of the same epitope, therefore, there are only 6 unique epitopes. <sup>c</sup> Calculated by dividing the MHCII affinity score of the highest scored epitope by the maximum predicted score for that haplotype. <sup>d</sup> Calculated by dividing the MHCII affinity score of the highest scored epitope by the affinity score of the 2<sup>nd</sup> highest scored unique epitope. <sup>e</sup> Not formally demonstrated but heavily suspected as one of the possible autoantigens involved in MS.

Our analysis shows that in all these diseases, the best ranked epitope for the MHC haplotype either monopolized the MHC presentation pathway or outmatched the second ranked by at least twofold, fulfilling the competitiveness clause for the establishment of RI.

Given their autoimmune origin, antigen availability (stoichiometry) is not a limiting factor and therefore, not an obstacle for the establishment of RI, which could become the ultimate driver of pathogenesis.

Taken together, analysis of these three autoimmune diseases indicates that the same RI mechanisms producing enhanced SELI in murine schistosomiasis, apply to human pathogenesis and show that RI can be a determinant of immunopathogenesis in CD4 T cell mediated diseases.

## Discussion

In the late 1800s, the advent of the paradigm of antigens as elicitors and steerers of the immune response inaugurated modern immunology, and a decades long effort to clarify how the former influence the latter. Unraveling this topic, though, proved itself so intricate that, to date, it has not yet been fully solved, still, some important fundamentals were established like: most immune responses target only a few dominant epitopes, determined by the interaction of three epitope-intrinsic properties: stoichiometry, affinity for MHC molecules and affinity for TCR, in a phenomenon called immunodominance (ID).

The first to recognize the importance of ID were the virologists who observed that minimal changes to an immunodominant epitope often turned self-limited viral diseases into chronic ones[26], however, similar evidence for CD4 mediated processes did not acknowledge ID an equivalent role in MHCII driven immune responses, still, over the years, the literature has accumulating examples where a single dominant epitope, presented to a particular CD4 clonotype, by a specific MHCII haplotype results in a disproportionately polarized response that resists immunomodulation and generates disease: murine leishmaniasis[8], experimental allergic encephalomyelitis[27], Lyme disease[23], respiratory syncytial virus pneumonitis[28] and schistosomiasis[17] among others. Our work aimed at deconstructing this process into its constituents to test whether this singular connection between a disproportionately dominant epitope and a specific MHCII haplotype we called RI, could answer for the pathogenic conversion of the CD4 response and we found that a) neutralizing RI abrogates disease, b) inducing RI enhances disease and c) antigens proven or suspected to generate human disease, fulfill the required conditions to produce RI.

These results suggest that RI, essentially an exceptional antigen presenting situation, could be the reason CD4 immunomodulation fails and becomes pathogenic, and correcting it could prevent/eliminate diseases like type I diabetes, pemphigus, celiac disease, multiple sclerosis, and others. According to this rationale, the only difference between schistosomiasis and all these diseases that result from a “rogue” CD4 response, is the location of the target epitope, respectively: liver, pancreas, skin, intestine, or brain. They also help explain the long known but never fully understood connection between MHC genes and autoimmune disease where disease would result from the domination of the MHC presentation pathway, by an inexhaustible antigen with disproportionate high affinity for MHC molecules, that creates such an intense and polarized CD4 response, that it cannot be modulated.

RI neutralization strategy may be useful even if the identity of the eliciting epitope is unknown since all one needs to counter it, is to prevent the installation of RI and all that is needed for it is the characterization of the MHC haplotype. One instance where this strategy could be helpful is the prevention of rejection in organ transplantation.

Mechanistically, one hypothesis to explain how RI neutralization works is that it would prime a naïve CD4 population into terminally differentiated Th2 egg-specific cells, which, upon the start of oviposition, migrates to the egg/granuloma interface and counteracts the polarizing effects of Sm-p40_234-246_RI. Supporting evidence for the role of *ad-hoc* pre-primed CD4 populations comes from the work of Julia et al., showing that the strong IL-4 response by Vβ4α8^+^ CD4s upon Leishmania infection actually originated from memory cells primed in the intestine by the microbiota[29].

In summary, our work shows that restrictive immunodominance may be a pathogenic determinant in CD4 mediated diseases, which may be manipulated for therapeutic purposes.

## Materials and methods

### Mice and Infection

CBA and SJL mice were acquired from University of Campinas animal facility and C57BL/6 mice were purchased from São Paulo Federal University. Mice were kept and monitored at our dedicated research animal facility.

*S. mansoni* cercariae were graciously provided by Dr. Pedro Paulo Chieffi of the Institute of tropical diseases from University of São Paulo.

For each experiment, female mice were infected at 6 weeks of age by intraperitoneal injection of 150 *S. mansoni* cercariae and euthanized 8 wpi.

### Morphometric and statistical analysis

Livers were collected in 10% formaldehyde and cut into 5μm thick, HE stained slides for analysis with an Eclipse50 microscope and digital image acquisition system (Nikon Instruments, Melville, NY). Only granulomas with a single visible central egg had their area measured with the NIS-Elements D 3.0 imaging software (Nikon Instruments). Statistical analysis consisted of the one-way ANOVA with Tukey’s multiple comparisons test (GraphPad Prism 6, La Jolla California).

### Cell culture, CD4 and APC purification and preparation

Immediately post euthanasia, mesenteric lymph nodes (MLN) were aseptically removed and manually dissociated into cell suspensions in cRPMI [RPMI1640 (Hyclone Laboratories, Logan, UT), supplemented with 10% FCS(Hyclone), 4 mM L-glutamine, 80 U/ml penicillin and 80 μg/ml streptomycin (Hyclone), 1 mM sodium pyruvate (Hyclone), 10 mM HEPES, 1x non-essential amino acids (Sigma, St. Louis, MO) and 6 ×10^−5^ M 2-ME]. This suspension was treated with Tris-ammonium chloride erythrocyte lysis buffer for 15 min, quenched with cRPMI, washed 3x, tested for cell concentration and viability and then diluted for CD4 magnetic purification according the protocol of CD4-T-cell-isolation kit (Miltenyi Biotech, Auburn, CA). APCs were prepared from the spleens of at least 2 uninfected mice by the same protocol and treated for 20 minutes at 37°C with mitomycin C 50μg/ml (Sigma).

### Epitope selection and synthesis

Evaluation and selection of haplotype specific dominant epitopes was performed *in silico* by submitting sequences of schistosome egg proteins published in ENTREZ and SchistoDB databases, for analysis with the RANKPEP algorithm[30].

Epitopes selected for the Sm-p40_234-246_ RI neutralization experiment were synthesized at Life Technologies (Grand Island, NY). CPP1380-1397, the RI inducing epitope for the H-2^b^ haplotype, was synthesized by Dr. Clovis Nakai at São Paulo Federal University, who also kindly provided random peptides to be used as controls and in mock cocktail experiments.

### Neutralization/induction of RI

For the neutralization of Sm-p40_234-246_ RI, CBA mice underwent subcutaneous immunization with a cocktail comprised of 2μM of each of the 4 competitor epitopes (Table 1) totalizing 8μM/mouse/immunization (Fig. 1), diluted in PBS prior to emulsification with adjuvant. The first immunization was performed 2wpi emulsified in CFA, the 2^nd^, at 4wpi with IFA. Induction of RI in C57BL/6 mice followed the same protocol but using 5uM of CPP1380-1397 instead.

### Cytokine analysis

For cytokine analysis, 1.5×10^6^ CD4s were incubated with 1.5×10^6^ syngeneic APCs at 37°C, 5% CO2 with or without addition of the appropriate antigens, or concanavalin A(Sigma) as a control. After 72 hours, supernatants were collected, filtered with a 0.2μm syringe filter (Millipore, Darmstadt, Germany) and analyzed with CBA analysis kit (BD, Carlsbad, CA), according to manufacturer’s protocol, using a FACSCalibur cytometer (BD).

## Author contributions

Eduardo Finger: conceived the project, wrote, and submitted grant proposal, assembled the laboratory, trained laboratory members, designed and performed experiments Thaissa Melo Galante Coimbra: performed experiments Alessandra Finardi de Souza: performed experiments

## Acknowledgments

Work funded and supported by FAPESP grant 06/02976-1

## Abbreviations

APC: Antigen presenting cell
CD4s: CD4-positive T cells
ID: Immunodominance
LD: Ligand density
MHC: Major Histocompatibility antigens
MHCII: MHC class II molecules
RI: Restrictive Immunodominance
SELI: Schistosome egg-induced liver immunopathology
TCR: T cell receptor
wpi: Weeks post infection

## Footnotes

* The situation where one epitope overwhelmingly dominates the MHCII presentation pathway thus producing an exceedingly high ligand density.

## References

1. Finger, E., Thermodynamics as the driving principle behind the immune system. Einstein (Sao Paulo), 2012. 10(3): p. 386–388.

2. Walsh, K.P. and K.H. Mills, Dendritic cells and other innate determinants of T helper cell polarisation. Trends Immunol, 2013. 34(11): p. 521–30.

3. Parish, C.R. and F.Y. Liew, Immune response to chemically modified flagellin. 3. Enhanced cell-mediated immunity during high and low zone antibody tolerance to flagellin. J Exp Med, 1972. 135(2): p. 298–311.

4. Murray, J.S., et al., MHC control of CD4+ T cell subset activation. J Exp Med, 1989. 170(6): p. 2135–40.

5. Tao, X., et al., Strength of TCR signal determines the costimulatory requirements for Th1 and Th2 CD4+ T cell differentiation. J Immunol, 1997. 159(12): p. 5956–63.

6. Kuchroo, V.K., et al., B7-1 and B7-2 costimulatory molecules activate differentially the Th1/Th2 developmental pathways: application to autoimmune disease therapy. Cell, 1995. 80(5): p. 707–18.

7. Hernandez, H.J., A.H. Sharpe, and M.J. Stadecker, Experimental murine schistosomiasis in the absence of B7 costimulatory molecules: reversal of elicited T cell cytokine profile and partial inhibition of egg granuloma formation. J Immunol, 1999. 162(5): p. 2884–9.

8. Launois, P., et al., IL-4 rapidly produced by V beta 4 V alpha 8 CD4+ T cells instructs Th2 development and susceptibility to Leishmania major in BALB/c mice. Immunity, 1997. 6(5): p. 541–9.

9. Hernandez, H.J., et al., Schistosoma mansoni: genetic restriction and cytokine profile of the CD4 + T helper cell response to dominant epitope peptide of major egg antigen Sm-p40. Exp Parasitol, 1998. 90(1): p. 122–30.

10. Sadler, C.H., et al., IL-10 is crucial for the transition from acute to chronic disease state during infection of mice with Schistosoma mansoni. Eur J Immunol, 2003. 33(4): p. 880–8.

11. Rutitzky, L.I., et al., Disruption of the ICOS-B7RP-1 costimulatory pathway leads to enhanced hepatic immunopathology and increased gamma interferon production by CD4 T cells in murine schistosomiasis. Infect Immun, 2003. 71(7): p. 4040–4.

12. Hernandez, H.J., et al., Expression of class II, but not class I, major histocompatibility complex molecules is required for granuloma formation in infection with Schistosoma mansoni. Eur J Immunol, 1997. 27(5): p. 1170–6.

13. Chikunguwo, S.M., et al., The cell-mediated response to schistosomal antigens at the clonal level. III. Identification of soluble egg antigens recognized by cloned specific granulomagenic murine CD4+ Th1-type lymphocytes. J Immunol, 1993. 150(4): p. 1413–21.

14. Hernandez, H.J. and M.J. Stadecker, Elucidation and role of critical residues of immunodominant peptide associated with T cell-mediated parasitic disease. J Immunol, 1999. 163(7): p. 3877–82.

15. Stadecker, M.J. and H.J. Hernandez, The immune response and immunopathology in infection with Schistosoma mansoni: a key role of major egg antigen Sm-p40. Parasite Immunol, 1998. 20(5): p. 217–21.

16. Stadecker, M.J., H.J. Hernandez, and H. Asahi, The identification and characterization of new immunogenic egg components: implications for evaluation and control of the immunopathogenic T cell response in schistosomiasis. Mem Inst Oswaldo Cruz, 2001. 96 Suppl: p. 29–33.

17. Finger, E., et al., Expansion of CD4 T cells expressing a highly restricted TCR structure specific for a single parasite epitope correlates with high pathology in murine schistosomiasis. Eur J Immunol, 2005. 35(9): p. 2659–69.

18. Pfeiffer, C., et al., Selective activation of Th1- and Th2-like cells in vivo--response to human collagen IV. Immunol Rev, 1991. 123: p. 65–84.

19. Murray, J.S., et al., Major histocompatibility complex (MHC) control of CD4 T cell subset activation. II. A single peptide induces either humoral or cell-mediated responses in mice of distinct MHC genotype. Eur J Immunol, 1992. 22(2): p. 559–65.

20. Murray, J.S., et al., Functional CD4 T cell subset interplay in an intact immune system. J Immunol, 1993. 150(10): p. 4270–6.

21. Kumar, V., et al., Major histocompatibility complex binding affinity of an antigenic determinant is crucial for the differential secretion of interleukin 4/5 or interferon gamma by T cells. Proc Natl Acad Sci U S A, 1995. 92(21): p. 9510–4.

22. Nelson, C.A., et al., A negatively charged anchor residue promotes high affinity binding to the MHC class II molecule I-Ak. J Immunol, 1996. 157(2): p. 755–62.

23. Gross, D.M., et al., Identification of LFA-1 as a candidate autoantigen in treatment-resistant Lyme arthritis. Science, 1998. 281(5377): p. 703–6.

24. Salentijn, E.M., et al., Celiac disease T-cell epitopes from gamma-gliadins: immunoreactivity depends on the genome of origin, transcript frequency, and flanking protein variation. BMC Genomics, 2012. 13: p. 277.

25. Riedhammer, C. and R. Weissert, Antigen Presentation, Autoantigens, and Immune Regulation in Multiple Sclerosis and Other Autoimmune Diseases. Front Immunol, 2015. 6: p. 322.

26. Pewe, L., S. Xue, and S. Perlman, Infection with cytotoxic T-lymphocyte escape mutants results in increased mortality and growth retardation in mice infected with a neurotropic coronavirus. J Virol, 1998. 72(7): p. 5912–8.

27. Waldner, H., et al., Fulminant spontaneous autoimmunity of the central nervous system in mice transgenic for the myelin proteolipid protein-specific T cell receptor. Proc Natl Acad Sci U S A, 2000. 97(7): p. 3412–7.

28. Varga, S.M., et al., Immunopathology in RSV infection is mediated by a discrete oligoclonal subset of antigen-specific CD4(+) T cells. Immunity, 2001. 15(4): p. 637–46.

29. Julia, V., et al., Priming by microbial antigens from the intestinal flora determines the ability of CD4+ T cells to rapidly secrete IL-4 in BALB/c mice infected with Leishmania major. J Immunol, 2000. 165(10): p. 5637–45.

30. Reche, P.A., et al., Enhancement to the RANKPEP resource for the prediction of peptide binding to MHC molecules using profiles. Immunogenetics, 2004. 56(6): p. 405–19.

